# Active sensing in groups: (what) do bats hear in the sonar cocktail party nightmare?

**DOI:** 10.1101/817734

**Authors:** Thejasvi Beleyur, Holger R. Goerlitz

**Affiliations:** Acoustic and Functional Ecology, Max Planck Institute for Ornithology, Seewiesen, Germany

**Keywords:** active sensing, bioacoustics, group behavior, psychoacoustics, sonar interference

## Abstract

Active sensing animals perceive their surroundings by emitting probes of energy and analyzing how the environment modulates these probes. However, the probes of conspecifics can jam active sensing, which should cause problems for groups of active sensing animals. This problem was termed the cocktail party nightmare for echolocating bats: as bats listen for the faint returning echoes of their loud calls, these echoes will be masked by the loud calls of other close-by bats. Despite this problem, many bats echolocate in groups and roost socially. Here, we present a biologically parametrized framework to quantify echo detection in groups. Incorporating known properties of echolocation, psychoacoustics, spatial acoustics and group flight, we quantify how well bats flying in groups can detect each other despite jamming. A focal bat in the center of a group can detect neighbors for group sizes of up to 100 bats. With increasing group size, fewer and only the closest and frontal neighbors are detected. Neighbor detection is improved for longer call intervals, shorter call durations, denser groups and more variable flight and sonar beam directions. Our results provide the first quantification of the sensory input of echolocating bats in collective group flight, such as mating swarms or emergences. Our results further generate predictions on the sensory strategies bats may use to reduce jamming in the cocktail party nightmare. Lastly, we suggest that the spatially limited sensory field of echolocators leads to limited interactions within a group, so that collective behavior is achieved by following only nearest neighbors.

**SIGNIFICANCE STATEMENT:** Close-by active sensing animals may interfere with each other. We investigated if and what many echolocators fly in a group hear – can they detect each other after all? We modelled acoustic and physical properties in group echolocation to quantify neighbor detection probability as group size increases. Echolocating bats can detect at least one of their closest neighbors per call up to group sizes of even 100 bats. Call parameters such as call rate and call duration play a strong role in how much echolocators in a group interfere with each other. Even when many bats fly together, they are indeed able to detect at least their nearest frontal neighbors – and this prevents them from colliding into one another.

## INTRODUCTION

Active sensing animals use self-generated energy to sense their surroundings by analyzing how objects around them change the emitted energy (1). Bats emit loud ultrasonic calls, and detect objects around them by listening to the echoes (2, 3) reflected off these objects. Active sensing is an effective sensory modality when the animal is solitary. However, when multiple active sensing animals emit pulses of energy in close proximity, they may ‘jam’ each other and mutually interfere with their ability to detect objects in their environment (1, 4). If groups of echolocating bats mutually jam or mask each other, they would not be able to detect each other. Due to the intense jamming, individuals would have a progressively difficult time detecting the echoes reflecting off their neighbors, and thus not detect them at all. Without detecting each other, groups of individuals cannot show collision free flight. However, many bat species are very gregarious, and fly and echolocate together in groups of tens to millions of bats. Bat groups also show coordinated behaviors in cave flights, evening emergences and mating swarms (5, 6). How is their ability to detect each other impaired with increasing group size? How many of its neighbors does a bat actually detect in the presence of intense jamming? What strategies may improve echo-detection and thus neighbor detection when many active sensing animals are together? We present biologically parametrized simulations to answer how bats manage to echolocate in the face of intense jamming.

In human psychophysics, the sensory challenge in perceiving an auditory cue among other similar sounds has been called the ‘cocktail party problem’ (7, 8). When applied to bat echolocation, the cocktail party ‘problem’ has been elevated to the ‘cocktail party nightmare’, given the repetition rate, similarity and high amplitude of echolocation calls. On top of these factors, is the non-linear increase in the number of masking sounds with increasing group size (9). Empirical studies to date have investigated the cocktail party nightmare from a sender’s perspective (sensu 7, 9). Through field observations, playback studies and on-body tags (11–22) we now know a range of echolocation strategies that bats show under challenging acoustic conditions. Bats can increase their call intensity, alter their call duration and frequency range, or suppress calling in the presence of conspecifics and noise playbacks (11, 20, 23, 24). In contrast to the many reports of bats’ response to noisy conditions-very little work has been done in conceptually understanding how receiver strategies might contribute to dealing with the cocktail party nightmare (25, 26). To our knowledge, biological modelling of the cocktail party nightmare from a receiver’s perspective that includes the details of bat echolocation and auditory processing is lacking. We fill this gap in conceptual understanding by presenting a biologically parametrized model based on the known properties of bat audition and the acoustics of a multi-bat echolocation scenario. We quantified how well a bat flying with conspecifics can perceive its neighbors in terms of the returning echoes it detects. Through our simulations we arrive at a sensory estimate of what a bat in the cocktail party nightmare may be detecting, if anything at all.

## MATERIAL AND METHODS

We model the echolocation of frequency-modulating (FM) bats. The calls of FM bats are typically downward frequency-modulated and of short duration (≤5 ms). Each call is followed by a longer silence (80-150 ms) called the interpulse interval (27). FM bats thus sense their world ‘stroboscopically’ by emitting a call and listening for the returning echoes in the interpulse interval (28). In the absence of any loud conspecific calls, a bat is able to hear all returning echoes and thus to detect all objects around it. However, in the presence of other loud bat calls, some of its own returning echoes may be masked. In that case, the bat will hear a few or none of the returning echoes. This corresponds to the bat detecting a few or none of the surrounding objects. In the cocktail party nightmare the ‘objects’ each bat is trying to detect are its neighbors.

Our model of the cocktail party nightmare is designed to describe the auditory scene (9) of a bat emerging from a cave in a group as it echolocates on the wing. A focal bat flying in a group of *N* bats may detect up to *N-1* of its neighbors (excluding itself), which is equivalent to hearing *N-1* returning echoes. The focal bat receives two kinds of loud masking sounds that interfere with the detection of its neighbors: 1) the *N-1* loud calls emitted by other bats in the group, and 2) the secondary echoes created by the call of a neighboring bat, reflecting once off another bat, and arriving at the focal bat. Every neighboring bat call generates *N-2* secondary echoes, meaning that the focal bat can receive up to *N-1xN-2* secondary echoes (**Fig. 1**). We implemented a spatially explicit 2-dimensional simulation of bat echolocation, sound propagation and sound reception and include mammalian auditory phenomena to quantify how many and which neighbors a bat can detect in the sonar cocktail party nightmare. We then explored how changes in group size and in sender strategies affect neighbor detection in a group.

**Fig. 1.**
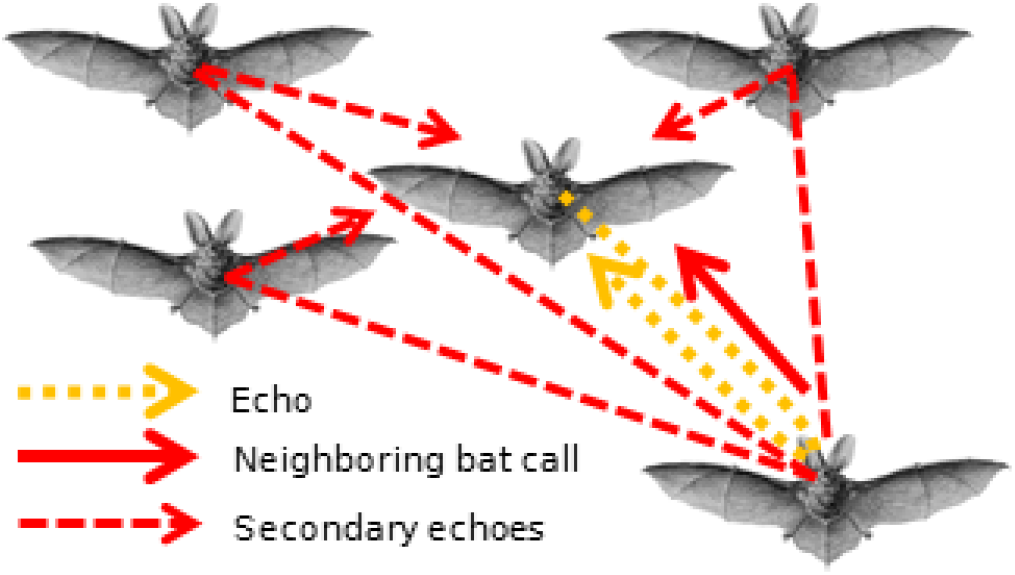
Schematic of the cocktail party nightmare. Arrows indicate the different types of sounds received by a focal bat: it needs to hear the echoes returning from its own calls (orange) to detect its neighbors, despite the masking by the calls of neighboring bats (solid red) and their secondary echoes (dashed red). Here, only one target echo off a single neighbor, only one representative neighboring bat call, and its set of secondary echoes are shown. In total, for a group of *N* bats, the focal bat will receive *N-1* echoes, *N-*1 neighboring bat calls and *N-1xN-2* secondary echoes. Bat drawing: *Kunstformen der Natur* (Ernst Haeckel, 1899).

### Model scenarios

We ran two model scenarios to test the effect of 1) increasing group size and of 2) variation in call parameters, group geometry and acoustic parameters on neighbor detection. In all models, we used the central-most bat in the group as the focal bat.

#### Scenario 1: Effect of group size on neighbor detection

We simulated groups of 5, 10, 30, 50, 75, 100 and 200 well-aligned bats with identical echolocation and hearing properties flying at a minimum inter-bat distance of 0.5 m (**Table 1** for full model parameters). The number and location of neighbors detected by the focal bat were recorded in every simulation run.

**Table 1.** Model parameters for both model scenarios. Scenario 1 modelled the effect of group size, while other parameters were fixed, resulting in 7 parameter combinations (one per group size). Scenario 2 modelled the effect of other relevant parameters, while group size was kept constant at 100 bats, resulting in a combined set of 1200 parameter combinations.

| *Parameter* | **Scenario 1: Effect of Group Size** | **Scenario 2: Effect of call parameters, group geometry and acoustics** |
| --- | --- | --- |
| *Group size* | 5, 10, 30, 50, 75, 100, 200 | 100 |
| *Interpulse interval (ms)* | 100 | 25, 50, 100, 200, 300 |
| *Call duration (ms)* | 2.5 | 1, 2.5 |
| *Source level (dB SPL re 20μPa at 1m)* | 100 | 94, 100, 106, 112, 120 |
| *Minimum inter-neighbor distance (m)* | 0.5 | 0.5, 1.0 |
| *Group heading variation (°)* | 10 | 10, 90 |
| *Atmospheric attenuation (dB/m)* | -1 | 0, -1, -2 |
| *Acoustic shadowing* | Yes | No, Yes |

#### Scenario 2: Effect of call parameters, group geometry and acoustic parameters on neighbor detection

Here, we varied other parameters relevant to the cocktail party nightmare (**Table 1**) while keeping group size constant (N=100, i.e., the largest group size from Scenario 1 with biologically relevant neighbor detection rate). We varied call parameters (interpulse interval, call duration, source level), group parameters (heading variation, minimum inter-bat spacing) and acoustic parameters (atmospheric absorption, acoustic shadowing).

### Model implementation

Each model run simulated one inter-pulse interval of the focal bat, and we calculated the timing and received level of all sounds (target echoes, masking calls, and secondary echoes) that arrived at the focal bat during that inter-pulse interval. Each model run simulated a series of sounds that arrived during an interpulse interval following the focal bats’ call, based on a spatially explicit distribution of a group of bats (**SI Appendix, Schematic S1**). At the beginning of every model run, *N* bats were placed in a 2D space with randomly assigned heading directions. For each neighboring bat, we calculated its angle and distance to the focal bat. The received level was calculated based on a common source level for all bats, spherical and atmospheric spreading over each call’s and echo’s travel distance, and acoustic shadowing. Acoustic shadowing is the reduction in received level of a sound due to obstructions in its path. A sound in the cocktail party nightmare may pass around obstacles (other bats) as it propagates from source to receive. The reduction in received level was measured and calculated as a linear function of the number of bats obstructing the path between source and receiver (See SI Section 1.9). For target and secondary echoes, we also considered monostatic and bistatic target strengths measured in this paper (see SI Section 1.8).

The arrival time of target echoes within the interpulse interval was determined according to the two-way travel time to the echo-reflecting neighboring bat. The arrival time of masking calls and secondary echoes was uniformly random within the interpulse interval. The random arrival time assignment of calls and secondary echoes recreates the non-coordinated echolocation of all bats in the group. It is unlikely that multiple bats in large groups can coordinate their calls effectively, and independent calling has been reported even in small groups of four bats (29).

All bats in a group were identical in their calling properties, and we treated all sounds as constant tones of equal duration, i.e., we did not explicitly model spectral emission, propagation and reception properties. The only difference between each of the sounds was their path and source of sound production. The omission of spectral properties is a conservative choice that assumes maximal masking of the primary echoes, thus allowing us to study the role of intensity differences and temporal separation between target echoes and masking sounds.

Once we calculated the timing and received level of all sounds at the focal bat, we accounted for directional hearing sensitivity (**SI Appendix, Fig. S3**) and spatial unmasking. Spatial unmasking describes the reduction in experienced masking as the arrival angle between masker and target sound increases (30, 31). We simulated spatial unmasking by the reduction of a masker’s effective received level based on its angular separation to an echo. For each echo, the same masker will have a different effective masking level as its relative angle of arrival will be unique for each echo. We thus calculated the effective masking level of each masker for each echo. The effective masking levels of all maskers were then combined to form a time-variant and echo-specific ‘masker SPL profile’ (**SI Appendix, Fig. S5D**). This is essentially the joint sound pressure level of all maskers over time. We then expressed this echo-specific masker SPL profile in relation to the echo’s SPL, thus obtaining a relative ‘echo-to-masker ratio profile’ (**SI Appendix, Fig. S5E**). This is equivalent to a signal-to-noise ratio profile, where the echo is the signal and the masker profile is the noise.

In addition to angular separation, signal detection is also determined by the temporal separation between signal (echo) and masker (24, 32, 33). Masking increases as the masker arrives closer in time to the echo. Masking occurs over longer durations when maskers arrive before the signal (forward masking) than afterwards (backward masking). We recreated the asymmetric masking by a ‘temporal masking envelope’ temporally centered at the echo (**SI Appendix, Fig. S1**). The echo was considered heard if the echo-to-masker ratio profile was above the temporal masking envelope. We allowed short drops of the echo-to-masker ratio profile below the temporal masking envelope, for a combined maximum duration of less than 25% of an echo’s duration (of 1 or 2.5 ms). Alternatively, we defined an echo to be masked (= not heard), if the echo-to-masker ratio profile was below the temporal masking envelope for more than 25% of the echo duration. The 25% threshold was an arbitrarily chosen conservative value to prevent rare bursts of high sound pressure level that are unlikely to affect echo detection biologically.

### Model parametrization

We implemented a detailed set of echolocation, group and sound properties in our model, including call and hearing directionality, spatial unmasking, temporal masking, group geometry and details of sound propagation. These properties were parameterized based on published results wherever available. Acoustic shadowing and target strengths (monostatic and bistatic) of bats were specifically measured for this work. All details of the model parameters including our respective measurements and on model implementation are presented in the Supplementary Information.

## RESULTS

### Effect of group size on neighbor detection

At group sizes of five and ten, the focal bat hears per call the echoes of most or all of its neighbors (median: 4 and 8 echoes at N=5 and 10, respectively; **Fig. 2**). At progressively larger group sizes, the median number of detected neighbors drops to between 4-0 at group sizes of 30-200. Yet even in a group of 100 bats, while the median number of detected neighbors is zero, the 90^th^ percentile is one, showing that a neighbor is not detected with each call, but occasionally. Beyond a group of 100 bats, the focal bat typically detects no neighbors at all. The initial rise in detected neighbors in groups of 5-30 bats is primarily caused by the increased number of neighbors that could be detected, which is soon counteracted by the intense masking that rises non-linearly with group size.

**Fig. 2.**
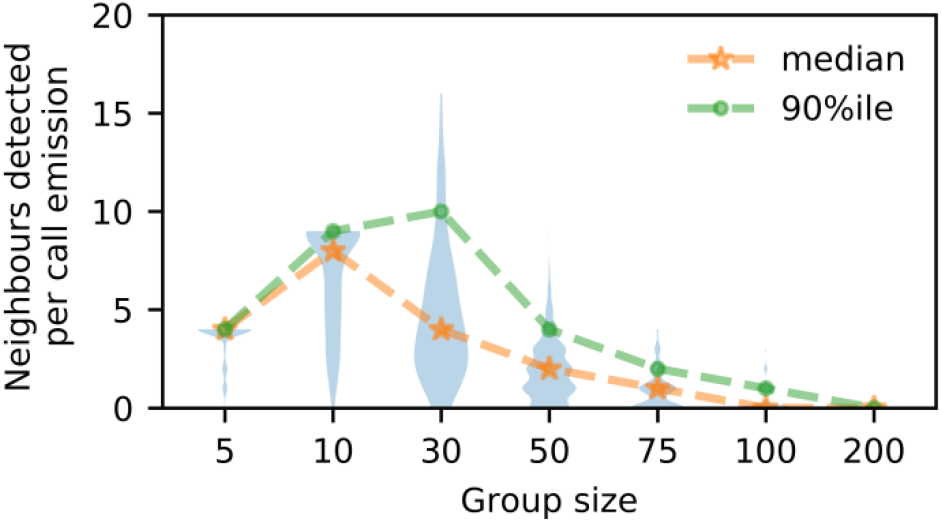
Number of detected neighbors per call by a focal bat in the center of a group. The initial rise in the number of detected neighbors is because there are indeed more neighbors and the degree of masking is negligible. However, with increasing group sizes, most of the neighbors cannot be detected any more, and progressively fewer neighbors are detected per call. Violin plots show the distribution of the number of neighbors detected per call, and their median and (stars, orange) and 90^th^ percentile (dots, green).

We next derived the probability of detecting at least one neighbor, which allows describing the average rate of neighbor detection (**Fig. 3A**, blue). At smaller group of 5 to 30 bats, the focal bat detects at least one neighbor per call at above 0.95 probability. At larger group sizes (50–100), the probability of detecting at least one neighbor drops rapidly to 0.3 per call in a group of 100 bats, and is basically zero for a group of 200 bats (0.004 probability). A bat (with 10 Hz calling rate) flying in a group of 100 bats will thus detect at least one neighbor around 3 times per second (~3 Hz detection rate), while a bat flying in a group of 30 bats will detect at least one neighbor almost every time (9.5 Hz detection rate). The probability of detecting multiple bats per call is lower than just detecting at least one bat (**Fig. 3A**). Yet, even in a group of 50 bats, the focal bat has a probability of detecting at least 2 and 4 neighbors per call of about 50 and 10%, respectively.

**Fig. 3.**
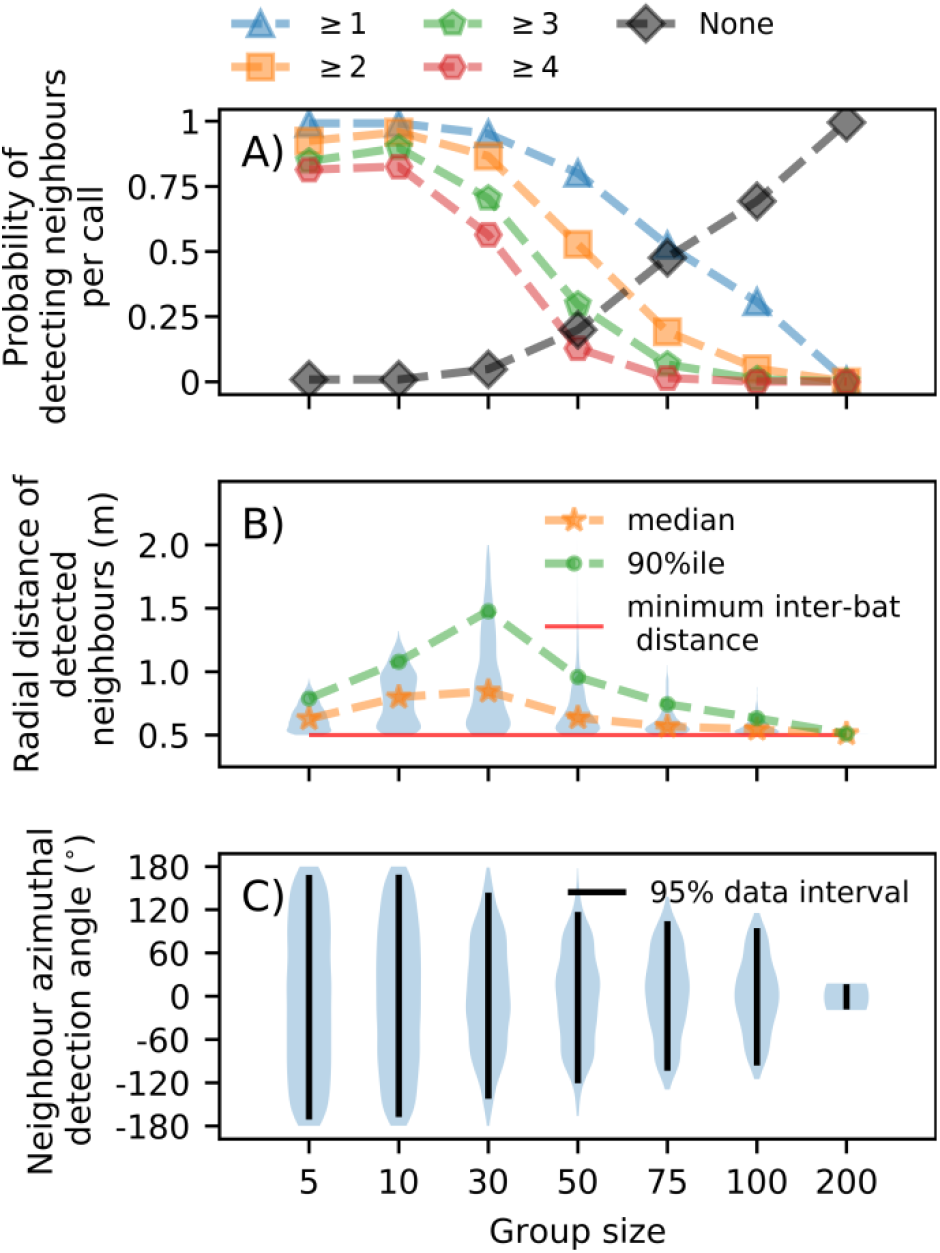
Characterization of the focal bat’s perception. **A)** The probability of detecting ≥*X* neighbors per call (X=1,2,3,4, or none). Even in groups of up to 100 bats, the focal bat has a ~0.3 probability of detecting at least one neighbor per call. In even larger groups (200 bats), no neighbors are detected anymore. **B)** With increasing group size, a focal bat only detects its closest neighbors. Initially, the radial distance of detected neighbors increases because the spatial extent of a group increases with group size (at 5, 10, 30 bats: radius = 0.75, 1.12; 1.97 m), but it then drops down to the nearest neighbors beyond 30 bats. **C)** The azimuthal location of detected neighbors, showing a increasing frontal bias with increasing group size. Although neighbors were uniformly distributed in azimuth, the frontal bias of call and hearing directionality means that frontal returning echoes are louder than peripheral ones.

We next quantified which neighbors the focal bat detects. Detection is generally limited to nearby neighbors (**Fig. 3B**) and, with increasing group size, to neighbors in front of the focal bat (**Fig. 3C**). At a group size of 30 bats, the focal bat occasionally detects neighbors that are up to 2 m away in radial distance, which is the furthest neighbor distance. With increasing group sizes, despite the group being more spread out, the focal bat can only detect its nearest neighbors (e.g. neighbors at ~0.5 m in a group of 200 bats; **Figure 3B**). In the azimuthal plane, at small group sizes the focal bat initially detects neighbors all around it (95%ile-neighbor detection angle >=237° for up to 50 bats; **Fig. 3C**). With increasing group size, a frontal bias in neighbor detection appears (95%-neighbor detection angle: 191-35° for 100 and 200 bats; **Fig. 3C**).

### Effect of call parameters, group geometry and acoustic phenomena on neighbor detection

We next analyzed how variation in call parameters, group structure, and acoustic parameters affected neighbor detection. We fixed the group size to 100, as at this size, the focal bat could typically detect at most one neighbor (90%ile, **Fig. 2**) at 0.3 probability (**Fig. 3A**) per call. We thus reduced the output of each simulation run to a binary neighbor detection score of 1 (detection) or 0 (no detection). We analyzed the effect of each parameter on neighbor detection with a logistic regression, treating all parameters as categorical and using their value in the previous model as reference (parameter range in **Table 1**).

The call parameters *interpulse interval* and *call duration* showed the strongest effect (**Fig. 4; SI Appendix, Table S2**). Increasing the interpulse interval from 100 ms to 200 and 300 ms increases neighbor detection probability by about 15 and 75 times, while reducing it to 50 ms lowers neighbor detection to 0.05 (**Fig. 4A**). Shortening call duration from 2.5 ms to 1 ms led to 35x higher neighbor detection (**Fig. 4B**). Call source level had no effect (**Fig. 4C**). Group geometry also influenced neighbor detection probability, but less than changing call parameters. Flying at larger interbat distances of 1.0 m leads to worse neighbor detection (odds-ratio: 0.31) compared to denser groups with 0.5 m interbat distance (**Fig. 4D**). Groups where individuals head in a generally common direction have worse neighbor detection than groups with variable heading (or echolocation beam) direction (odds-ratio: 1.32, **Fig. 4E**).

Among the physical parameters, *acoustic shadowing* increased neighbor detection (odds ratio: 0.75) compared to simulations without acoustic shadowing, while *atmospheric attenuation* had a negligible effect (**Fig. 4 F,G**).

**Fig. 4.**
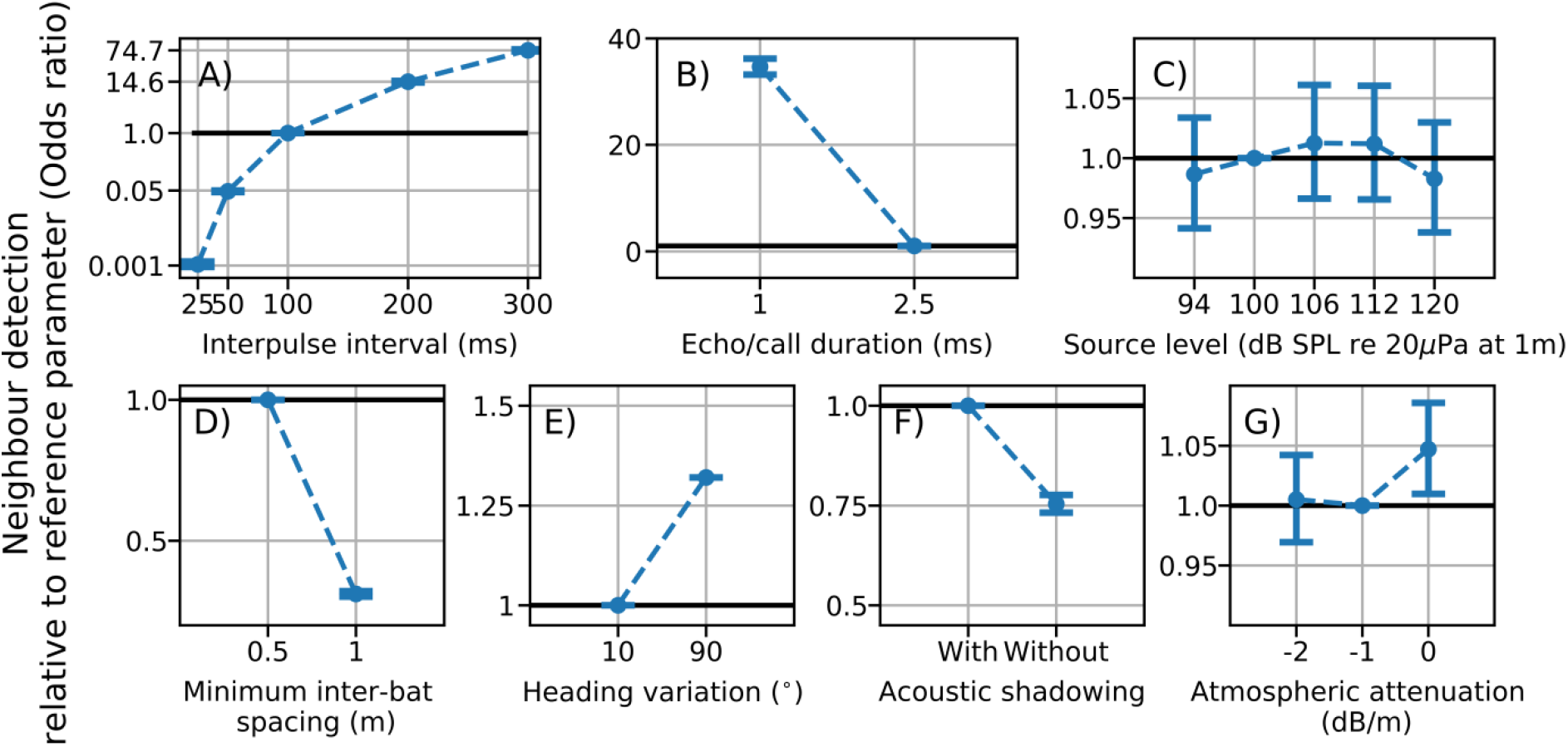
Effect of call parameters (A-C), group geometry (D-E) and acoustic parameters (F-G) on neighbor detection. Each plot shows the probability of neighbor detection (model estimate and 95% confidence interval of odds ratio) when changing model parameters relative to the reference parameter used in the previous simulations of scenario 1 (**Table 1**). Odds ratios above and below one indicate a higher and lower neighbor detection probability, respectively, indicated by the horizontal reference line. **A-C) Call parameters**: Longer interpulse intervals (A) and shorter call durations (B) increase neighbor detection probability, while call source level (C) has no effect. **D,E) group geometry**: Neighbor detection is better in groups that are tightly packed (D) and with higher heading variation (E). **F,G) Effect of acoustic parameters:** Acoustic shadowing by bats in groups improves neighbor detection probability (F), while atmospheric attenuation has a negligible effect (G).

## DISCUSSION

We present a conceptual framework to quantify what a focal bat experiences in the sonar cocktail party nightmare. We quantified the probability of detecting neighbors across a range of group sizes, which allows calculating the rate at which a focal bat detects its neighbors. When flying alone, a focal bat will detect objects around it at a rate equal to its call rate, while in a group, its object detection rate is reduced due to masking. We show that even in a group of 100 bats, bats still detect at least one neighbor per call about 3 times per second (for a 10 Hz call rate), while in smaller group sizes, neighbor detection rate is larger at 5-10 Hz. Bat echolocation is generally ‘stroboscopic’, meaning that information is received intermittently with time gaps (3). We suggest that bats in smaller group sizes still experience a sufficiently high information update rate for performing collision avoidance and neighbor following. With increasing group size, perception might become ‘hyper-stroboscopic’, i.e., so scarce that different sensorimotor heuristics might be required to maintain group coordination.

The low level of masking at smaller group sizes allows the focal bat to detect all its neighbors per call. With increasing group size, however, the focal bat detects maximally one neighbor per call in a group of 100 bats. This neighbor detection rate of at least one neighbor per call even in large group sizes provides a formal sensory basis for group movement in active sensing animals. While a bat in a large group cannot track the position of all its neighbors, it still can track the movement of a few neighbors, specifically those close to and in front of it. This reduction in rate, range and direction of detected neighbors has predictive consequences for the kind of collective behavior bat groups may show in nature. Many models of collective movement assume that each individual in a group detects the position and orientation of neighbors in the whole of its sensory volume, and then performs an ‘averaging’ across all neighbors to decide its next movement (34–37), leading to the impressive coordinated behaviors of fish schools and insect swarms (38, 39). As the number of neighbors that an individual detects decreases, more ‘limited interactions’ begin to dominate, causing anisotropy in the group structure (40, 41). For bats in the cocktail party nightmare, we predict that large groups may show higher anisotropy than smaller groups due to the limited number of neighbors that they can detect and react to. All things being equal, we predict that in large groups (>50 bats), the neighbors in the frontal field of a bat will have a disproportionate influence on its movement decisions. Bats in larger groups may thus maintain higher alignment with their frontal neighbors compared to bats in smaller groups.

Our simulations allow for a direct quantitative comparison of the effects of echolocation, group geometry and acoustic phenomena in group echolocation. Among the call parameters tested, reducing call rate (increasing interpulse interval) was most effective in increasing neighbor detection in jamming conditions, matching experimental evidence for reduced calling rate in *Tadarida brasiliensis* (19) (20). In contrast, other FM bat species increase their call rates in groups and background noise (11, 15, 42, 43). Likewise, our result that shorter call duration should improve neighbor detection is opposite to experiments showing that most bat species increased call duration in the presence of maskers (11, 23, 24, 43, 44), except (42). Lastly, our result of no effect of changing source level on neighbor detection might also seem to differ from experimental data showing that bats in laboratory conditions do increase source level in the presence of maskers (11, 23, 43, 44). While there might be species-specific differences, we suggest that these differences are mostly due to differences in experimental situations. Bats in these experiments experienced constant maskers, thus calling more often, for longer and for louder improved the bats’ signal redundancy, echo-to-masker ratio, and overall echo detection. In contrast, our model simulates group flight of many bats with simultaneous and uniform changes in their call parameters. When all bats in a group shorten call duration, this reduces the overall duration of masking sounds, thus improving echo detection. Likewise, when all bats in a group increase their call amplitudes to optimize their own echo-to-masker ratios, all bats will eventually call at their maximum, with no overall effect on neighbor detection. Analyzing bat calls in mass emergences is technically challenging and it remains unknown whether *T. brasiliensis* and other gregarious bat species reduce their call rate in the field.

Bat aggregations show a variety of structure across behavioral contexts, from well-aligned almost parallel flight during roost emergences, to more variable and less-aligned flight in mating swarms and when circling in limited cave volumes. We show that this group structure itself affects how well bats can detect each other. Bats detect their neighbors better in less-aligned groups compared to more aligned groups. During aligned emergence flight, the focal bat always receives loud frontally directed masking calls from bats behind it, in addition to the relatively loud side-calls emitted by neighbors to its left and right. In contrast, during less-aligned swarming flight, the relative orientation of the bats is more distributed and changing, with the focal bat experiencing a wider dynamic range of masker levels (i.e., louder and fainter masking calls originating from a wider range of directions around it). This increased dynamic masker range allows for better neighbor-echo detection, as there will be drops in echo-to-masker ratios due to changing received masker level. This effect is beneficial for enabling swarming flight, as the collision risk in less-aligned flight is likely higher compared to the more aligned emergence flight. Inter-individual distance is another parameter of group structure, and we show that neighbor detection is better in dense groups. This might seem unexpected given that the received SPL of the maskers is higher the closer the bats are. However, received echo levels are also higher when bats are closely spaced. Since echo SPL drops with 12 dB per doubling of distance, but masker call SPL only by 6 dB doubling of distance, the echo-to-masker ratio is higher at shorter than longer interbat distances. It would be interesting to examine if perhaps large groups in the field actually fly closer to each other than smaller groups.

While we only modeled neighbor detection for the central-most bat in a group, its position in the group (e.g., central, frontal or at the back) is likely to also have an effect on the number and received level of maskers, and thus on the number of detected echoes. However, we expect the obtained trends to remain qualitatively the same regardless of focal bat position. Particularly, we assume that masking will increase with group size, and only the exact group size at which a given level of masking (e.g. X% neighbor detection probability) is obtained will change depending on the focal bat’s position in the group.

We furthermore show that it is important to consider bats not only as sources of echoes to be detected and of masking sounds, but also as obstructions to sound that actually alleviate the cocktail party nightmare. While the detected echoes originate from nearby bats, they are typically not shadowed. In contrast, the masking calls and secondary echoes can arrive from distant neighbors, thus passing through multiple other bats. Shadowing thus consists of the overall reduction in masker levels, which increases echo-to-masker ratios for the comparatively loud echoes returning from nearby neighbors.

Our results show that the cocktail party may not be as much of a ‘nightmare’ as previously thought (9). We show that the modelled psychoacoustic, spatial and acoustic properties act together to alleviate the ‘nightmare’ into a ‘challenge’. When bats are flying in a multi-echo environment, our results show that a bat will always hear some echoes after a call emission, and very rarely no echoes at all. This parallels the phenomenon of auditory ‘glimpsing’ reported in the human auditory cocktail party where individuals may follow conversations by perceiving parts of detected speech rather than whole sounds (45).

### Improved echo-detection in real-world situations

We present a first order approximation to the sonar cocktail party nightmare, including many relevant biological, physical and auditory mechanisms. Bats are expert echolocators and can detect echoes and fly under challenging conditions (24, 46–48). Bats rapidly adjust their call behavior in terms of their call duration, source level and interpulse intervals (49, 50), integrate echoic information over multiple call emissions (51) and actively track objects by aiming their calls at them (52, 53). While we tested a range of different echolocation call parameters, our model implemented these parameters as fixed values that do not vary over time, thus lacking the dynamic nature of a real bat in the field.

Furthermore, we did not model the spectral content of echo or masker sounds, and analyzed echo detection based on a fixed threshold of echo-to-masker-ratio. In contrast, real echolocation calls possess a time-variant spectral pattern that is species and even individual-specific (13, 54), which can reduce echo masking. Masking is strongest when target and masker overlap both in time and in frequency (i.e., fall within the same ‘critical band’ of the auditory system, (32, 55). The frequency-modulation of bat calls means that even when maskers and echoes partially overlap in time, they will not necessarily overlap in frequency, thus reducing the likelihood of masking. The individuality of bat calls may help a bat reject the secondary echoes from other bats’ calls by forming separate auditory streams (56) for its own echoes and others’ echoes. Given the scarcity of empirical data to parametrize the effect of spectral differences on echo detection in masking conditions, we did not include it in the model, thus simulating a conservative worst case scenario where all sounds lie in the same frequency band. Additionally, attentional processes strongly improve target detection by improving the required signal-to-noise ratio despite the presence of maskers with similar time-frequency structure (57). Under real-world conditions, it is likely that masking in groups is even less than simulated here.

Due to the scarcity of published data, the inter-individual and inter-specific variation in the temporal and spatial masking functions used in our model is unknown. The temporal masking envelope will arguably be similar in many bat species, showing the typical mammalian pattern of increased target detection threshold with reduced temporal separation between target and masker. Spatial unmasking occurs through the nonlinear interaction of pinnae shape, cochlear and higher auditory processing (30, 58). As pinna shape and associated acoustic receiver characteristics strongly vary in echolocating bats (59), leading to species-specific spatial unmasking and echo detection rates in the cocktail party nightmare.

## CONCLUSION

We provide a conceptual framework to explain how active sensing animals such as echolocating bats successfully navigate in groups despite mutually jamming each other. The intense jamming in groups might lead to individuals only detecting their nearest frontal neighbors, which might drive limited interactions within a group. We also show that call parameters and group geometry determine the challenge in the cocktail party nightmare. Recent advances in on-body acoustic tags (42, 60), signal analysis (61) and acoustic tracking (62) of echolocating animals in the field might facilitate future experimental validation of our model predictions. As our model formulation is not constrained to echolocation in bats, it can be parametrized to other echolocators such as oilbirds, swiftlets and odontocetes (63, 64) that also echolocate in groups and suffer from cocktail-party like conditions.

## Supporting information

Beleyur_Goerlitz2019_SI

## Author Contributions

Conceptualization: HRG, TB; Code: TB; Analysis and Visualization: TB; Writing: TB, HRG

## Acknowledgements

We thank the members of the Acoustic and Functional Ecology Group for insightful comments and support, Claire Guérin for contributing to code development, the MPI for Ornithology for excellent research infrastructure, and two anonymous reviewers for helpful comments.

## Funding

TB was funded by a doctoral fellowship from the German Academic Exchange Service (DAAD) and the IMPRS for Organismal Biology. HRG was funded by the Emmy Noether program of the German Research Foundation (DFG, GO 2091/2-1, GO 2091/2-2). The simulation runs in this paper were funded by a Google (Research Credits) grant to TB.

## Data and code availability

All code required to replicate the simulations, results, analyses and figures in this paper are available at this Zenodo repository link: https://doi.org/10.5281/zenodo.3514156. All raw data and code required to replicate the results of the experimental parametrizations (target strength and acoustic shadowing) are available at the following repository link: https://doi.org/10.5281/zenodo.3469845.

## Notes

Declarations of interest: none

#### Summary of Updates

updated SI

https://doi.org/10.5281/zenodo.3514156

