## Supplementary material for "Active sensing in groups: (what) do bats hear in the sonar cocktail party nightmare?": Beleyur_Goerlitz2019_SI

#### Modelling active sensing reveals echo detection even in large groups of bats

**This PDF file includes:**

Supplementary methods
Figures S1 to S5
Tables S1
Schematic S1
SI References

### SUPPLEMENTARY METHODS

#### 1. Model parametrization

Here, we present the details of how we parametrized our model of the sonar cocktail party nightmare, based on empirical data of behavioral studies in bats and our own measurements.

##### 1.1 Temporal masking envelope

We derived the echo-to-masker sound pressure level (SPL) ratios for forward, backward and simultaneous masking from two empirical target-detection studies in echolocating bats (**Table S1**). We only chose studies where the target and masker were co-located along the same direction. Both studies presented the ratio between the echo and masker SPL at target detection for various delays between echo and masker arrival time. We linearly interpolated (in a piecewise fashion) the echo-to-masker SPL ratios between each of the time delays measured in the studies to obtain the full temporal masking envelope ranging from -0.65 and +24 ms delay of the target echo relative to the masker edge (**Fig. S1**).

##### 1.2 Spatial unmasking function

Sümer et al., 2009 (1) performed a backward masking study to address spatial unmasking in the bat *Eptesicus fuscus*. In a two-alternative forced-choice paradigm, they increased the angular separation between a masker object and a target object while also varying the level of the target echo (by varying object size and thus object target strength).

We define the spatial unmasking function as the echo-to-masker SPL ratio at just-noticeable echo detection as a function of angular separation. To obtain the echo-to-masker level ratios, we subtracted the object target strengths from the masker object target strength. We normalized the spatial unmasking function to the co-localised echo-masker case (i.e., when both echo and masker arrive from the same direction). This describes the reduction in echo-to-masker ratio required for echo detection as a function of angular separation between target and masker, compared to the co-localized case. We digitized the data points from Fig. 4B of Sümer et al. (2009) by hand with WebPlotDigitiser (2) to obtain the target's target strength as a function of angular separation. Masker target strength was given by Sümer et al. (2009) as -14.5 dB. We then calculated the target-masker SPL ratios and interpolated them with a quadratic polynomial fit (**Fig S2**). The interpolated data was then further upsampled to 0.47° intervals. As Sümer et al. (2009) only measured angular separations up to 23°. We conservatively used the echo-masker SPL ratio at 23° also for all larger angular separations.

##### 1.3 Call and hearing directionality

Echolocation calls have a directional beam shape, meaning that the emitted SPL generally decreases with increasing angular distance from the main call direction, which has the highest call SPL (**Fig. S3**). The highest call SPL is typically towards the front, and reduces towards the back of the bat. Despite this directionality and additional variation with call frequency and behavioral context (3–6), bats still emit a significant amount of sound pressure into the backward direction. The average call SPL behind a bat is about 14 dB lower than in the forward direction (6, 7). Call directionality leads to a drop in the effective number of masking calls from neighbors as only those calls arriving from a limited range of directions will have sufficiently high SPL (**Fig. S3**). For example, in an emergence situation with approximately parallel flight directions, the focal bat will receive the loudest calls from those bats flying behind it. Similarly, the lowest received call levels in an emergence will be from those bats flying in front of the focal bats.

Like call production, hearing is also directional. The pinna structure of a bat attenuates or amplifies the same sound depending on its direction of arrival (8, 9). In *Myotis daubentonii*, hearing directionality between 35-45 kHz leads to an average amplification of frontally arriving sounds by 4 dB in comparison to those arriving from behind. We used data of Giuggioli et al (2015) to describe the average call and hearing directionality of our modelled bats (**Fig. S3**).

##### 1.4 Atmospheric attenuation

Ultrasound in air is heavily attenuated by atmospheric attenuation, even over short distances of a few meters (10, 11). Atmospheric attenuation will reduce the received SPL of a masker or echo at the focal bat. We chose a range of values for the atmospheric attenuation coefficient $\alpha$ between 0 to -2 dB/m. These values approximate the atmospheric attenuation experienced by a bat calling at very low (≤20 kHz, ~0 dB/m) to high (60 kHz, ~-2 dB/m) peak frequencies.

##### 1.5 Geometric attenuation

Sound pressure level reduces with increasing distance from the source, called geometric attenuation. For all sounds in our model (target echoes, masking calls and secondary echoes), we implemented spherical geometric spreading, i.e., uniform spreading of sound in all directions (12).

##### 1.6 Group geometry

A group of bats might organize themselves tightly or well spread in the field. The spacing between bats will decide how loud the returning target echoes, masking calls and secondary echoes are. We simulated a group of bats by placing individual bats on a 2D plane using the Poisson disk algorithm (13). The advantage of using the Poisson disk algorithm is that points are spaced relatively uniformly in space compared to a random placement of points from two independent distributions. The other advantage of the Poisson disk algorithm is that it allows the specification of a minimum distance between two points. For the first simulations varying group size only, we chose 0.5 m as inter-bat distance (see Table 1, main text), matching the average interbat-distance in dense swarms of *T. brasiliensis* in the field (14). In addition to 0.5 m minimum inter-bat distance, we also studied how a sparser 1.0 m minimum inter-bat distance affects neighbor detection (see Table 1, main text). The Poisson disk arranged bats showed average inter-bat distances of between 1-1.5 times the specified minimum distance between points.

##### 1.7 Heading variation

Active sensing animals are known to 'scan' their environments by emitting energy in varying directions of interest according to the behavioral context (15, 16). Bats alter the shape and direction of their sonar beam while they fly (3, 17, 18). The directions into which each bat in a group aims its calls could affect how well each bat in the group can detect echoes. A group of bats calling into the same direction may experience high masking, as a focal individual will receive many loud calls from the bats behind it. In contrast, a group of bats calling into a larger range of directions may experience less masking. The focal bat may receive a mix of fainter off-axis calls and loud on-axis calls from neighbors.

We simulated the scanning behavior of individual bats in the group by setting a heading angle for each individual. Each individual called into the direction of its heading angle, and we chose two levels of variation of heading angles in the group. Groups with a low heading variation were all pointing their beams in more or less the same direction. Groups with high heading variation were pointing their beams in a wider range of directions. A low heading variation simulates an emergence situation where each bat is calling approximately into the overall flight direction of the group. A high heading variation simulates a swarming situation where each bat is calling at a unique direction. Given the lack of empirical data to guide our estimates, we chose ±10° for the low heading variation, and ±90° for the high heading variation. The heading angle for each individual was randomly drawn from a uniform distribution covering the respective range.

**1.8 Monostatic and bistatic target strength of a flying bat**

Quantifying the received levels of echoes and secondary echoes requires knowledge of the target strength of a bat when emitter and receiver are at the same and at different locations. Here, we measured monostatic and bistatic target strengths (19) of a flying stuffed *Myotis myotis* bat. Monostatic target strengths refer to the situation where the emitter and receiver are at the same location, i.e., they are the same bat (this is the 'classical' target strength usually considered in echolocation research). Bistatic target strength refers to a situation where the emitter and receiver are at different locations, i.e., the receiving bat hears the echo of a call that was emitted by another bat, i.e. a secondary echo.

In the simulations, all incoming and outgoing sounds at the bat are between abs(0-180)°. Sounds with 0° angle are along the heading direction of the focal bat. Sounds arriving/reflecting on the left have negative angles ($0 \geq \theta \geq -180°$), and those on the right have positive angles ($0 \leq \theta \leq 180°$).

**Methods:** We ensonified a stuffed *Myotis myotis* with outstretched wings, which was suspended from the ceiling at ~1 m height and placed on a rotating base, which could be rotated in 45° steps. A speaker (electrostatic Polaroid, custom built) and microphone (CM16/CMPA, Avisoft Bioacoustics, Glienicke, Germany) were placed at a 1 m radial distance to the center of the bat (**Fig. S4**). The speaker emitted linear frequency modulated sweeps between 96-20 kHz, with durations of 170 µs, 1 ms and 2 ms at 92 dB rms SPL re 20 µPa at 1 m. The speaker was driven by a custom-built amplifier with input from a soundcard (Player 216H, Avisoft Bioacoustics, 1 MHz sampling rate). The microphone signal was recorded simultaneously with an attenuated version of the speaker signal on a multichannel soundcard (USG 416H, Avisoft Bioacoustics, 500 kHz sampling rate). The microphone had a noise floor of 24 dB rms SPL re 20 µPa. All echoes were recorded at ≥22dB signal-to-noise ratio. The experiment was performed in the middle of a large empty room (~4x4x2 m) to temporally separate bat echoes from background echoes.

We ensonified the bat from front (0°) to back (180°) in steps of 45°. We assumed that the bat was symmetrical and thus did not ensonify angles from 180-360°. The angular separation between the speaker and the microphone was also altered in steps of 45° between -180° to +180°. This resulted in 40 target strength measurements (5 sound directions x 8 angular separations).

The integrated target strength (20) of the recordings were calculated by subtracting the energy of recordings with the bat from those without the bat at the expected time window of echo arrival. The echo level was calculated in rms by taking the square root of the energy.

**Results:** The monostatic target strength of a flying stuffed *Myotis myotis* bat at various orientations was between -43 and -34 dB at 1 m distance, matching the general range of previously published values (21). The bistatic target strength, which was used to calculate the received level of the secondary echoes, was between -44 and -10 dB across all combinations of emitter-receiver locations.

For further details on experimental protocol, raw data and reproduction of generated results, please refer to the archived Jupyter notebooks at this link: https://zenodo.org/record/3469845#.Xa4QEugzZIA.

##### 1.9 Acoustic shadowing in bat groups with varying number of bats and inter-bat spacing

As multiple bats fly together in a group, the bats themselves will block all sounds travelling between an emitter and a receiver. Essentially, the bats themselves act as obstacles that cause acoustic shadowing, reducing the received sound pressure level at the focal bat. In a large group, multiple bats may shadow a sound as it moves from the emitter and to the receiver. We quantified acoustic shadowing in a series of playback experiments that varied the inter-bat spacing (0.5 and 1.0 m) and the number of bats (1 – 6) in a line.

**Methods:** A microphone (CM16/CMPA, Avisoft Bioacoustics) and speaker (Polaroid, custom-built) were placed at a fixed distance of 9.9 m apart, facing each other. We hung 1 to 6 "model bats" made of foam with paper wings at 0.5 or 1.0 m distance to each other from a string running above the speaker to the microphone. The designed model bat showed acoustic shadowing similar to that of the stuffed *Myotis myotis* used in the target strength measurements described in **1.8**.

The speaker was placed as far as possible from the microphone to calculate acoustic shadowing without the effects of speaker directionality. The speaker played back a variety of 7 ms Tukey windowed signals consisting of pure tones (20, 35, 50, 100 kHz) and a downward modulated linear sweep (100-15 kHz). Each signal type was played back 15 times at ~4% duty cycle. Multiple signal types were used to obtain a generalized estimate of shadowing across a wide range of call peak frequencies and call types. The playback signals and recordings are available here: https://doi.org/10.5281/zenodo.3469845. Additionally, we also recorded the same playback without model bats being present. We calculated acoustic shadowing as the reduction in received level by subtracting the received level (in dB rms) without bats from the received level with bats. We performed a linear regression of attenuation as a function of factors *number of bats* and *inter-bat-distance*, to estimate the amount of acoustic shadowing caused per bat and the spacing between them.

**Results:** Bats effectively shadowed the sound, with strong effects of the inter-bat-spacing and the number of bats. Bats at 0.5 m distance in front of the receiving bat (=microphone) reduced the received SPL by 5.17 dB (SEM=0.44, t=-11.639, 95% CI =-6.05,-4.30), while bats at 1.0 m interbat-spacing reduced the received SPL by 1.85 dB (SEM=0.44, t=-4.164, 95% CI =-2.72,-0.98). Each bat reduced received SPL by 0.83 dB (SEM=0.08, t=-9.852, 95% CI =-0.99,-0.66).

For further details on experimental protocol, raw data and reproduction of generated results, please refer to the archived Jupyter notebooks at this link: https://zenodo.org/record/3469845#.Xa39_ugzZIA.

#### 2. Model implementation

Here, we present how we implemented the parameters described before into our final model, and how our model was initialized and run.

##### 2.1 Model idea

The idea of our model is to analyze the relative timing and sound pressure level of target echoes, masking calls and secondary echoes at a focal bat flying in a group of other bats. Each model iteration thus analyzed one single interpulse interval, i.e., the time after emission of one call until the emission of the next call by the focal bat. Within that interpulse interval, the focal bat received the echoes from its own call that reflected off the neighboring bats, the calls of those neighboring bats, and the secondary echoes which originate from the calls of the neighboring bats reflecting off other neighboring bats (Figure 1, main text).

We placed groups of bats in a 2D plane with various inter-bat distances and heading directions. We then calculated received timing and SPL of all sounds based on realistic assumptions about call properties, mammalian auditory characteristics and sound physics.

All echoes, calls and secondary echoes were considered to be equal in duration, amplitude envelope, and frequency composition. Frequency composition was not explicitly specified, which is a conservative modelling choice that maximizes masking potential (22) and makes our model generalizable to multiple bat species. All sounds were treated as having a constant amplitude envelope (i.e., no amplitude modulations), but they differed in the sound pressure level received by the focal bat.

##### 2.2 Model initialization

Each model iteration consisted of distributing bats in a 2D plane and assigning each bat a heading direction. This spatial distribution was used to calculate the arrival times and received level of target echoes, masker calls and secondary echoes within the interpulse interval (**Fig. 1**). The interpulse interval was discretized into time bins of 1 μs duration. Each received target echo corresponded to one neighbor. The arrival time of each echo was calculated using twice the distance between focal and neighboring bat.

The arrival time of masker calls and secondary echoes were chosen randomly. The random arrival time assignment of masker calls and secondary echoes is supported by the finding that groups of *Miniopterus fuliginosus* (23) do not coordinate their calling behavior, and seem to echolocate independently. Moreover, at large group sizes beyond a few bats it is unlikely that bats could effectively co-ordinate their call emission times.

**2.3 Target echo properties**

Target echoes are the echoes that the focal bat receives in response to its own echolocation call. In our model, the target echoes are echoes reflected off the neighboring bats. When a focal bat hears a target echo it means it has detected the corresponding neighboring bat.

An echo was defined as a sound occupying a block of time within the interpulse interval (**Fig. S5A**). Echoes were simulated to arrive at delays corresponding to the distance to the neighboring bat they reflected off, e.g. if a neighboring bat was at 1 m distance to the focal bat, then its echo arrived at a delay of 6.06 ms (at 330 m/s sound propagation). The received level of the returning echo was calculated based on emitted call source level into the direction of the neighboring bat, our monostatic target strength measurements of a bat, and geometric attenuation over the sound travel distance. If acoustic shadowing and atmospheric absorption were included in a simulation run, the received level was reduced based on the number of bats in the path and the atmospheric attenuation for the overall distance travelled by the echo. Echo arrival direction was determined based on the position of each neighboring bat.

##### 2.4 Masker call properties

Masker calls arrived at random time points with uniform probability within the interpulse interval (**Fig. S5,B**), based on the observed lack of call synchronization in groups of *Miniopterus fuliginosus* (23). Call directionality was based on the directionality function in Giuggioli et al., 2015, who fit a cosine based function to describe the overall call directionality of *Myotis daubentonii* echolocating in the field. We set the asymmetry parameter A to 7.0. We calculated the angle of call emission towards the focal bat for each conspecific bat based on its angular position (heading) and distance. We then calculated the effective source level into the direction of the focal bat by reducing the call's on-axis source level (**Table 1 in main text**) according to the call directionality function and the focal bat's relative position to the conspecific (**Fig. S3**). This reduced level was the final received level of the conspecific masker call. If acoustic shadowing and atmospheric attenuation were included in a simulation run, the received level was reduced based on the number of bats in the path and the overall distance travelled by the call.

##### 2.5 Secondary echo properties

Like the masking calls, secondary echoes arrived randomly with uniform probability in the interpulse interval (**Fig. S5C**). The received level of a secondary echo was based on the emitted call source level into the direction of the neighboring bat, our bistatic target strength measurements of a bat, and geometric attenuation over the sound travel distance. If acoustic shadowing and atmospheric absorption were included in a simulation run, the received level was reduced based on the number of bats in the path and the overall distance travelled by the secondary echo.

##### 2.6 Obtaining the masker sound pressure level profile

All sounds were treated as having a fixed received level (no envelope modulations). For each target echo, we calculated its unique masker SPL profile based on the relative timing, relative arrival directions and received levels of all masking calls and secondary echoes. This masker SPL profile was different for each target echo because the temporal and spatial properties of the masking sounds differ for each echo, resulting in different received levels and spatial unmasking (see main text for details). We first calculated the effective masker SPL for each masking sound by correcting for spatial unmasking based on the angular separation between the echo and the masker. All effective masker SPLs of all masking sounds together over time represent the complete masker sound pressure profile for each target echo (**Fig. S5,E**). When two or more maskers overlapped in time, we added their linear sound pressures to obtain their joint masking SPL. This approach assumes that overlapping maskers are coherent sound sources that constructively interfere. This is a conservative assumption that will maximize masking.

**2.7 Determining neighbor detection: the temporal masking envelope**

The masker SPL profile for each echo describes the received masking SPL over time. From the masker SPL profile, we created an echo-to-masker ratio profile by normalizing the SPL of the target echo to the masker SPL profile:

$$echo\text{-}to\text{-}masker\ \ ratio\ profile\ (dB) \\ = echo\ level\ (dB\ SPL) - masker\ sound\ pressure\ profile\ (dB\ SPL)$$

The echo-to-masker-ratio profile is comparable to a signal-to-noise-ratio: at 0 dB, echo and masker have the same SPL. The masker is louder than the echo for negative values, and the echo is louder than the masker for positive values.

To determine whether a given target echo was heard or not, we compared the echo-to-masker ratio for this echo with the temporal masking envelope (see 1.1). The temporal masking envelope describes the echo-to-masker ratio at which masking occurs as a function of relative timing between echo and masker. Using the temporal masking envelope is important because masking does not only occur when the masker coincides with the  echo, but also when the masker does not overlap with the echo and arrives before (forward masking) or after (backward masking) it. In our case, echo and masker had durations of only 1-2.5 ms, while a masker arriving at up to ~25 ms before and up to ~1 ms after the echo still causes some amount of masking. Thus, our temporal masker envelope had a duration of either ~27 or ~28.5 ms (**Fig. S5F**). We compared the echo-to-masker ratio profile to the temporal masking envelope. The echo was considered not heard if the echo-to-masker ratio profile lay below the temporal masking envelope, i.e., the echo-to-masker ratio was lower than required for echo detection. Alternatively, the echo was considered heard if the echo-to-masker ratio profile lay above the temporal masking function, i.e., the echo-to-masker SPL ratio was higher than required for echo detection.

However, as the echo-to-masker ratio continuously fluctuates over time, it is possible that it is not fully above or below the temporal masking envelope throughout the envelope's duration. We thus defined an echo to be masked (= not heard), if it was masked for more than 25% of its duration (of 1 or 2.5 ms). To calculate the total duration of masking, we analyzed the total duration that the echo-to-masker ratio was below the temporal masking envelope.. As long as the total masking duration was shorter than 25% of the echo duration (of 1-2.5 ms) the echo was considered detected. If this duration was longer than 25% of the echo duration, the echo was masked and the corresponding neighboring bat was considered not detected. This 25% threshold was set to make the simulated auditory system immune to short spikes in masking sound pressure level occurring during the temporal masking function.

#### 3. Open-source software used in the research

All simulation code, experimental data and results were made possible through the use of the NumPy (24), SciPy (25), Pandas (26), Matplotlib (27), Statsmodels (28), sounddevice (29), Anaconda (30) and CPython (31) open-source projects.

#### 4. Acknowledgements

We thank Renate Heckel and Felix Hartl for building the ensonification setup, Magnus Wahlberg for helpful discussions on the ensonifications, Henrik Brumm for permission to use Raum 1.03 for the ensonification experiments, and Mihai Valcu for facilitating simulation runs on the in-house server facility.

*Soc Am* 124(4):2123–2132.

10. Goerlitz HR (2018) Weather conditions determine attenuation and speed of sound : Environmental limitations for monitoring and analyzing bat echolocation. *Ecol Evol* 8(March):5090–5100.
11. Lawrence BD, Simmons JA (1982) Measurements of atmospheric attenuation at ultrasonic frequencies and the significance for echolocation by bats. *J Acoust Soc Am* 71(3):585–590.
12. Speaks CE (1996) *Introduction to Sound: Acoustics for the Hearing and Speech Sciences* (Singular Publishing Group, Inc., San Diego, California). Second.
13. Bridson R (2007) Fast Poisson disk sampling in arbitrary dimensions. *ACM SIGGRAPH 2007 sketches - SIGGRAPH '07*:22-es.
14. Theriault DH, et al. (2010) Reconstruction and Analysis of 3D Trajectories of Brazilian Free-tailed Bats in Flight. *Bost Univ Comput Sci Techinical Rep* (Figure 2):1–7.
15. Wisniewska DM, et al. (2015) Range-dependent flexibility in the acoustic field of view of echolocatingporpoises (Phocoena phocoena): Short title: Flexible field of view in echolocating porpoises. *Elife* 2015(4):1–29.
16. Bullock TH, Hopkins CD, Popper AN, Fay RR eds. (2005) *Electroreception* (2005 Springer Science+Business Media, Inc, New York).
17. Ghose K (2006) Steering by Hearing: A Bat's Acoustic Gaze Is Linked to Its Flight Motor Output by a Delayed, Adaptive Linear Law. *J Neurosci* 26(6):1704–1710.
18. Lee WJ, et al. (2017) Tongue-driven sonar beam steering by a lingual-echolocating fruit bat. *PLoS Biol* 15(12):1–15.
19. Richards MA, Scheer J., Holm WA (2010) *Principles of Modern Radar* (SciTech Publishing, Edison, New Jersey).
20. Cook JC (1985) Target Strength and Echo Structure. *Adaptive Methods in Underwater Acoustics*, ed Urban HG (D. Reidel Publishing Company), pp 155–172.
21. Götze S, Koblitz JC, Denzinger A, Schnitzler HU (2016) No evidence for spectral jamming avoidance in echolocation behavior of foraging pipistrelle bats. *Sci Rep*

6(July):1–13.

22. Yost WA (2007) *Fundamentals of hearing: an introduction* (Academic Press, London).
23. Hase K, et al. (2018) Bats enhance their call identities to solve the cocktail party problem. *Commun Biol* 1(1):39.
24. Oliphant TE (2006) *A guide to NumPy* (Trelgol Publishing USA).
25. Virtanen P, et al. (2019) SciPy 1.0--Fundamental Algorithms for Scientific Computing in Python. *arXiv Prepr*:1–22.
26. McKinney W (2010) Data Structures for Statistical Computing in Python. *Proc 9th Python Sci Conf (SciPy 2010)*.
27. Hunter JD (2007) Matplotlib: A 2D graphics environment. *Comput Sci Eng* 9(3):90–95.
28. Seabold S, Perktold J (2010) Econometric and statistical modeling with Python. *Proc 9th Python Sci Conf (SciPy 2010)*.
29. Geier M (2015) sounddevice 0.3.5. Available at: https://python-sounddevice.readthedocs.io/en/0.3.5/_modules/sounddevice.html.
30. Anaconda (2016) Anaconda Software Distribution, Vers 2-2.4.0, Nov. 2016, https://anaconda.com.
31. Van Rossum G (1991) Interactively Testing Remote Servers Using the Python Programming Language. *CWI Q* 4(December):283–303.
32. Siewert I, Schillinger T, Schmidt S (2004) Forward masking and its consequences for echo perception in the gleaning bat, Megaderma lyra (Echolocation in bats and dolphins. University of Chicago Press, Chicago).

**Schematic 1**: Pseudo-code of the steps in a simulation run to determine the detected neighbors per call emission.

1. Place *N* bats in group with minimum inter-bat distance
2. Choose bat closest to the center of the group as the focal bat
3. Populate interpulse intervals with maskers and echoes:
    a. Propagate maskers (calls and secondary echoes) and calculate their received levels according to the position and orientation of the source neighbors. Assign random timing within interpulse interval.
    b. Propagate echoes from focal bats' own call and calculate their arrival time and received levels according to the position and orientation of the neighbors.
4. Implement hearing directionality of the focal bat: amplify the received level of all sounds according to their relative angle of arrival
5. Per echo, determine if it was heard:
    a. Implement spatial unmasking by reducing the effective received level of all masking sounds based on their angular separation to the echo
    b. Combine all maskers over time to form a 'masker profile'
    c. Calculate the echo-to-masker profile', with reference to the echo level
    d. Implement temporal masking by checking if the relative echo-masker profile lies below the temporal masking envelope centered on the echo's location in the interpulse interval.

**Table S1:** Target-detection studies in echolocating bats used to extract echo-masker SPL ratio for our model. The time delay is the time between the edge of a masker and the target echo. A positive time delay indicates forward masking (masker arrives before the target), a negative time delay indicates backward masking (masker arrives after the target).

| Publication | Study species | Time delay (ms) | Masking condition | Echo-masker SPL ratio (dB) |
|---|---|---|---|---|
| Siewert et al. (2000) (32) | *Megaderma lyra* | 3 | Forward | -17 |
| | | 6 | Forward | -23 |
| | | 12 | Forward | -29 |
| | | 24 | Forward | -34 |
| Sümer et al. (2009) (1) | *Eptesicus fuscus* | -0.65 | Backward | -22.3 |

**Table S2: Results of the logistic regression** to quantify the effect of different parameter values on the odds ratio to detect at least one neighbor. Odds ratio values >1 indicate a higher probability of neighbor detection, while odds ratios <1 indicate a lower probability of neighbor detection.

| Parameter | Value tested | Reference value | Odds Ratio | Odds ratio-2.5 CI | Odds ratio-97.5 CI | Log Odds ratio | Log Odds ratio SEM | Z (log odds ratio estimate) | P > \|z\| |
|---|---|---|---|---|---|---|---|---|---|
| **Intercept** | | | 0.32 | 0.31 | 0.35 | -1.11 | 0.027 | -40.51 | 0.0 |
| **Heading variation (°)** | ±90 | ±10 | 1.32 | 1.28 | 1.36 | 0.28 | 0.015 | 18.35 | 0.0 |
| **Acoustic shadowing** | Yes | no | 0.75 | 0.73 | 0.78 | -0.28 | 0.015 | -18.63 | 0.0 |
| **Interpulse interval (ms)** | 25 | 100 | 0.001 | 0.0009 | 0.001 | -6.84 | 0.073 | -93.52 | 0.0 |
| | 50 | 100 | 0.048 | 0.046 | 0.050 | -3.04 | 0.023 | -134.14 | 0.0 |
| | 200 | 100 | 14.60 | 13.995 | 15.228 | 2.68 | 0.022 | 124.46 | 0.0 |
| | 300 | 100 | 74.68 | 70.497 | 79.122 | 4.31 | 0.029 | 146.49 | 0.0 |
| **Minimum interbat distance (m)** | 1.0 | 0.5 | 0.31 | 0.301 | 0.321 | -1.17 | 0.016 | -72.83 | 0.0 |
| **Sound duration (ms)** | 1 | 2.5 | 34.66 | 33.172 | 36.206 | 3.55 | 0.022 | 158.83 | 0.0 |
| **Source Level (dB SPL re 20 µPa at 1m)** | 94 | 100 | 0.99 | 0.941 | 1.034 | -0.01 | 0.024 | -0.57 | 0.57 |
| | 106 | 100 | 1.01 | 0.966 | 1.061 | 0.01 | 0.024 | 0.52 | 0.60 |
| | 112 | 100 | 1.01 | 0.966 | 1.061 | 0.01 | 0.024 | 0.50 | 0.62 |
| | 120 | 100 | 0.98 | 0.938 | 1.030 | -0.02 | 0.024 | 0.73 | 0.47 |
| **Atmospheric attenuation (dB/m)** | 0 | -1 | 1.05 | 1.010 | 1.086 | 0.05 | 0.019 | 2.48 | 0.01 |
| | -2 | -1 | 1.01 | 0.970 | 1.042 | 0.01 | 0.018 | 0.27 | 0.78 |

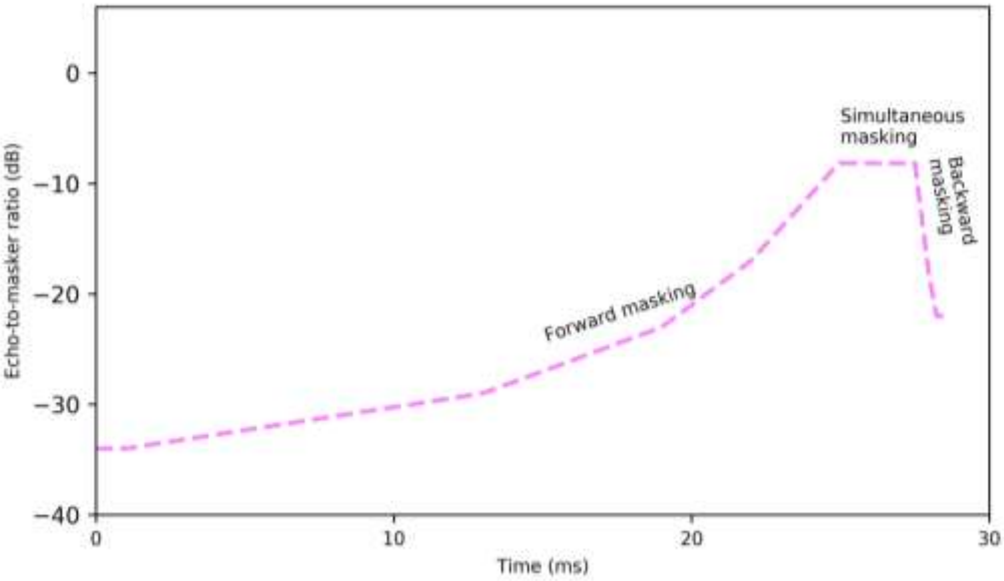


**Figure S1: The 'temporal masking envelope'** used to simulate temporal masking. The envelope represents the lower echo-to-masker ratios at which a bat can detect echoes for various echo-masker delays. The envelope is the equivalent of the lowest signal-to-noise ratios at which echoes can be detected over different time delays. The envelope is centered on the position of the echo, and has a long forward masking section (at times prior to the echo), and a short backward masking section (at times after the echo). The simultaneous masking region is equal to the length of the echo itself. If the echo-to-masker ratio profile is above the temporal masking envelope for most of its duration (i.e., the echo-to-masker SPL ratio was higher than required for echo detection), we considered an echo to be heard. If the echo-to-masker ratio profile is below the envelope for more than 25% of the echo's duration, the echo was considered not heard. Here the temporal masking envelope is shown for a 2.5 ms echo. Data and sources used to construct the temporal masking envelope are given in Table S1.

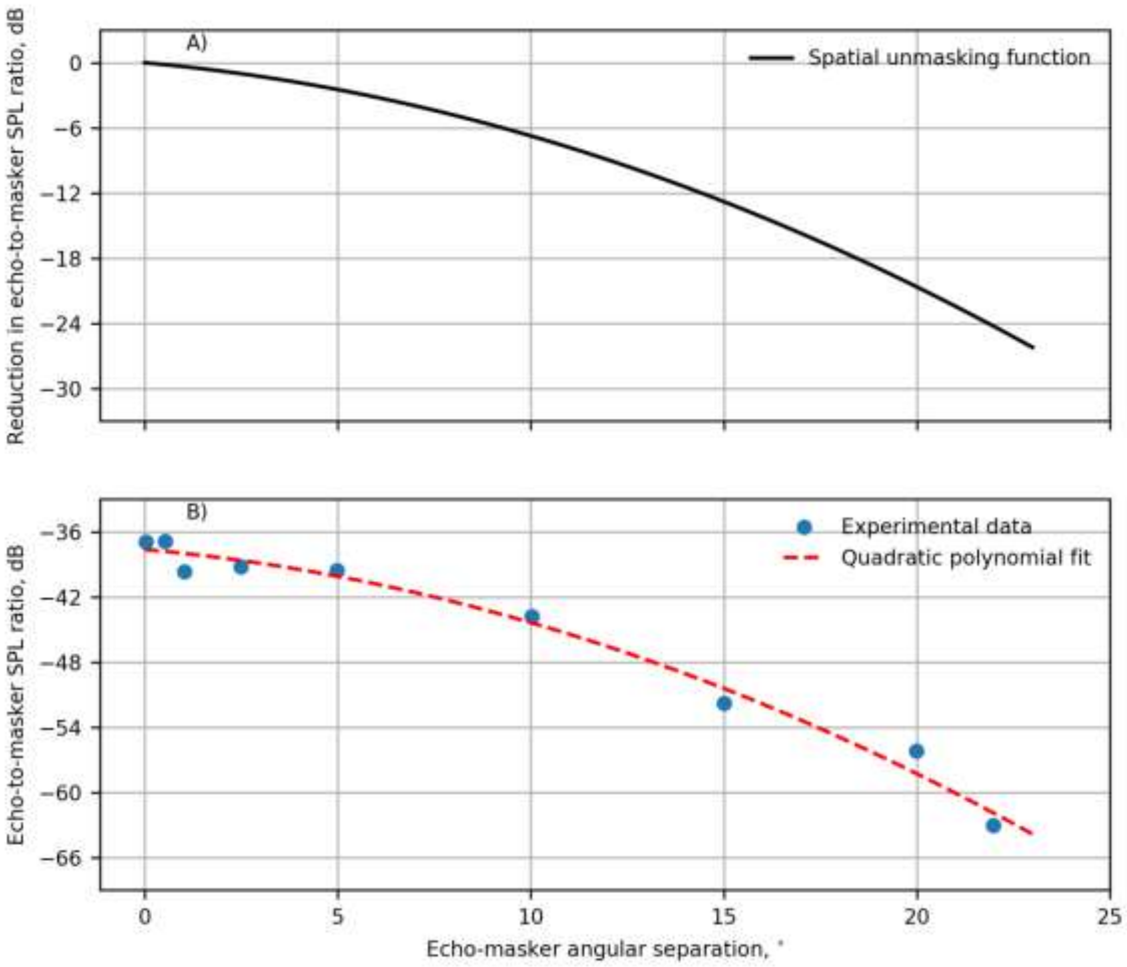


**Figure S2: Deriving the spatial unmasking function.** A) The 'spatial unmasking function' describes the reduction in echo-to-masker SPL ratio at echo detection as a function of angular separation between echo and masker. B) The original data set of Sümer et al. (2009) (blue dots) and our digitized and interpolated dataset (dashed red line). The error between the data and our interpolation is less than 2 dB. The spatial unmasking function (A) used in the simulations was obtained by normalising the echo-to-masker SPL ratio at zero degrees angular separation. This allowed the creation of a function which showed the reduction in required echo-to-masker SPL ratio with 0 dB reduction when echo and masker were co-located and greater reduction with angular separation.

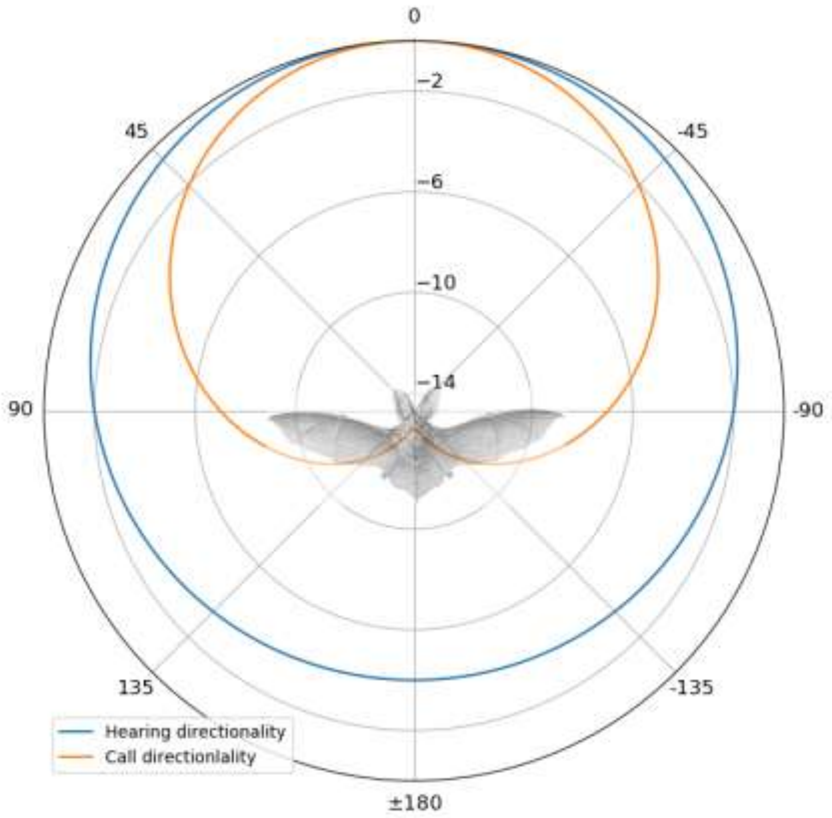


**Figure S3: Calling and hearing directionality of bats.** Call directionality (orange) is directional, with a difference of up to -14 dB in source level from front to back. Calls emitted to the front of a bat result in higher received levels of calls, echoes and secondary echoes. Hearing directionality (blue) is less directional, with a difference of up to -4 dB from front to back. Hearing directionality causes sounds arriving from the back to be perceived fainter than sounds arriving from the front. Bat drawing from *Kunstformen der Natur* (Ernst Haeckel, 1899).

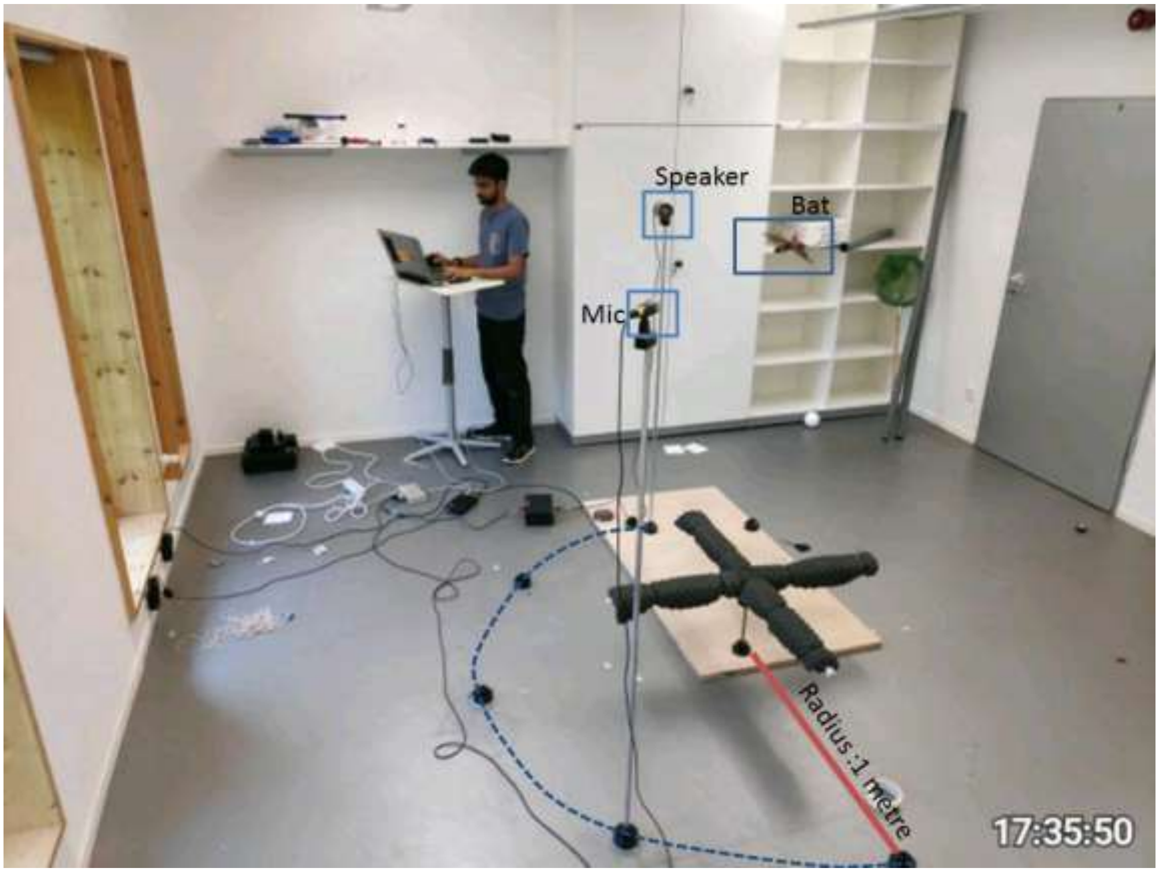


**Figure S4: The ensonification setup used to measure the monostatic and bistatic target strength of a bat as an acoustic target.** A stuffed *Myotis myotis* bat was hung at the same height as the speaker and microphone. The bat could be rotated in the azimuth. The microphone and speaker were placed at 1 m radius around the bat at various positions (black rounded plastic molds on floor) with a separation of 45° from each other. By a combination of bat orientation, microphone and speaker positions all possible incoming and outgoing relative angles were measured. Here the positions of the speaker and microphone for a bistatic target strength measurement with 135° angle between microphone and speaker are shown.

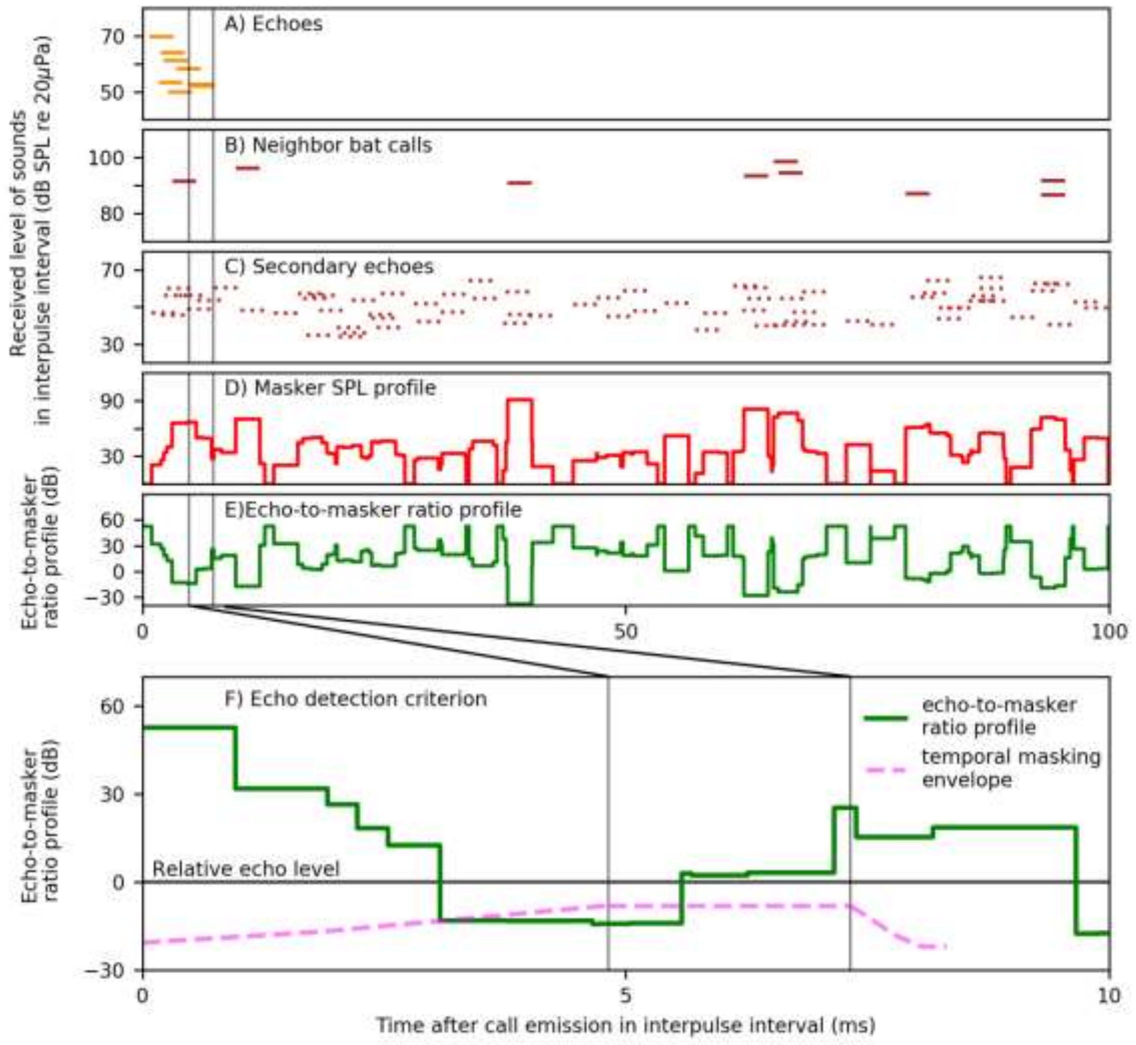


**Figure S5: Schematic representation of the sounds arriving in the interpulse interval for a simulation with a group of 10 bats (A-C) and how echo detection was determined (D-F).** A-E show the full interpulse interval of 100 ms, while F shows an enlargement of the first 10 ms.

**A)** Timing and received SPL of the individual echoes reflected off neighbors. The echoes arrive at delays corresponding to the neighbors' distance from the focal bat. The vertical lines single out one specific target echo to illustrate the simulated auditory system (see F).

**B)** Timing and received SPL of the calls from neighboring bats arriving randomly with a uniform probability over the interpulse interval.

**C)** Timing and received SPL of the secondary echoes in the interpulse interval, arriving randomly with uniform probability over the interpulse interval.

**D)** The masker SPL profile obtained by adding the effective masker SPL of all maskers (calls and secondary echoes) over time for the chosen echo. The effective masker SPL is the received SPL corrected for spatial unmasking based on the angular separation between the echo and the masker, using the spatial unmasking function in Fig. S2.

**E)** The echo-to-masker ratio profile obtained by normalizing the echo SPL to the masker SPL profile. 0 dB is the echo's relative SPL.

**F)** Determining whether an echo was heard or not, by comparing the echo-to-masker ratio profile (solid green) to the temporal masking envelope (dashed pink). If the echo-to-masker ratio is above the temporal masking envelope, then the echo was not masked. In contrast, if the echo-to-masker ratio is cumulatively below the temporal masking envelope for more than 25% of the echo's duration (of 1 or 2.5 ms), then the echo was considered masked. The vertical lines indicate the actual temporal location of the example echo from A). The temporal masking envelope is centered on the chosen echo. Here, the echo-to-masker ratio is below the temporal masking envelope for almost a whole echo duration, meaning that this echo was masked.
